# L2G: Repurposing Language Models for Genomics Tasks

**DOI:** 10.1101/2024.12.09.627422

**Authors:** Wenduo Cheng, Junhong Shen, Mikhail Khodak, Jian Ma, Ameet Talwalkar

## Abstract

Pre-trained language models have transformed the field of natural language processing (NLP), and their success has inspired efforts in genomics to develop domain-specific foundation models (FMs). However, creating high-quality genomic FMs from scratch is resource-intensive, requiring significant computational power and high-quality pre-training data. The success of large language models (LLMs) in NLP has largely been driven by industrial-scale efforts leveraging vast, diverse corpora and massive computing infrastructure. In this work, we aim to bypass the data and computational bottlenecks of creating genomic FMs from scratch and instead propose repurposing existing LLMs for genomics tasks. Inspired by the recently observed ‘cross-modal transfer’ phenomenon – where transformers pre-trained on natural language can generalize to other modalities – we introduce L2G, which adapts a pre-trained LLM architecture for genomics using neural architecture search (NAS) and a novel three-stage training procedure. Remarkably, without requiring extensive pre-training on DNA sequence data, L2G achieves superior performance to fine-tuned genomic FMs and task-specific models on more than half of tasks across multiple genomics benchmarks. In an enhancer activity prediction task, L2G further demonstrates its capacity to identify significant transcription factor motifs. Our work not only highlights the generalizability and efficacy of language models in out-of-domain tasks such as genomics, but also opens new avenues for more efficient and less resource-intensive methodologies in genomic research.

## Introduction

In recent years, large-scale pre-trained models, often referred to as foundation models (FMs) [1], have revolutionized the field of natural language processing (NLP). Models such as BERT [2], LLaMA [3], and GPT [4] leverage self-supervised pre-training on vast amounts of unlabeled text data to develop a rich and nuanced understanding of language. Similarly, DNA sequences can be viewed as strings of nucleotides, with recurring patterns analogous to reusable elements in natural language. This parallel has inspired the development of genomic FMs, such as DNABERT [5], Nucleotide Transformer [6], and HyenaDNA [7], which are pre-trained on large-scale genomic sequence data with subsequent fine-tuning. These models have shown potential in predicting complex genomic features, such as genomic element, chromatin state, and genome function.

However, constructing these domain-specific FMs from scratch is costly and resource-intensive. For instance, training the Nucleotide Transformer required approximately 174 billion tokens and 28 days of continuous training on 128 NVIDIA A100 GPUs. Even smaller models like DNABERT-2 required two weeks of training on 8 GTX 2080 Ti GPUs [8]. More recent architectures with fewer parameters than transformer-based models, such as HyenaDNA [7] and Mamba [9], still require substantial compute budget and massive training corpora (see **Fig**. 1B and **Table** S1 for details).

**Figure 1:**
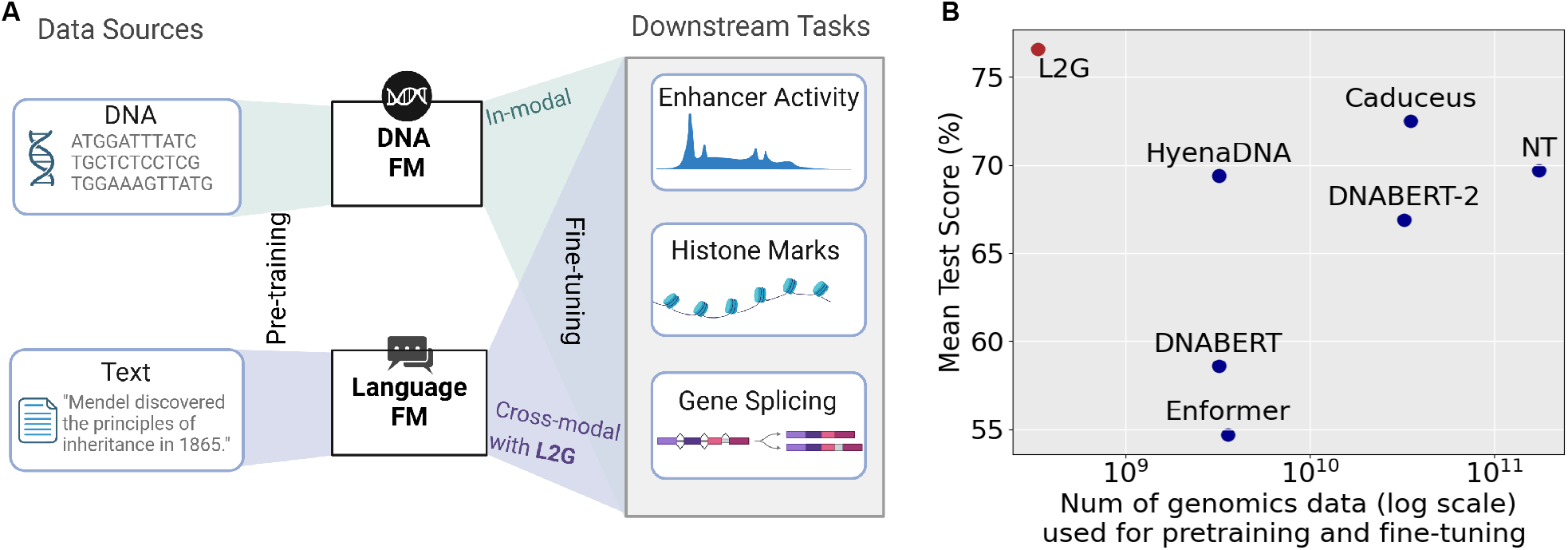
L2G is a data-efficient cross-modal fine-tuning method for genomics. **A**. Schematic overview contrasting genomic foundation models (FMs) with L2G during pre-training and fine-tuning. Genomic FMs pretrain with massive DNA sequencing data, while L2G bypasses genomic-specific pre-training altogether by leveraging existing pre-trained language models. Both approaches perform fine-tuning for specific downstream genomic task, but L2G’s three-stage cross-modal fine-tuning workflow on par with vanilla fine-tuning in terms of compute and data requirements. **B**. L2G (red dot) achieves a higher mean test score than leading genomic FMs on the Nucleotide Transformer benchmark, with higher values indicating better performance. By skipping pre-training, L2G requires significantly less genomic data and computational resources.

To address these challenges, we explore an alternative strategy to bypass genomic pre-training altogether. Our framework, Language-to-Genome (L2G), adapts pre-trained language models to genomic prediction tasks. This work is motivated by recent advances in the *cross-modal transfer* paradigm, which leverages the general reasoning capacity of pre-trained LLMs in domains such as protein property prediction [10] and solving partial differential equations [11]. These studies demonstrate that transferring existing models from well-studied text and vision modalities to scientific applications holds the promise of not only drastically reducing the required computational and data resources associated with pre-training, but also improving downstream model performance. L2G builds on a general-purpose cross-modal transfer approach [12] but incorporates neural architecture search and a novel three-stage training procedure (**Fig**. 2) to adapt to the unique features of genomics data, significantly enhancing empirical effectiveness. **Fig**. 1A contrasts L2G with the traditional approach of building genomic FMs.

**Figure 2:**
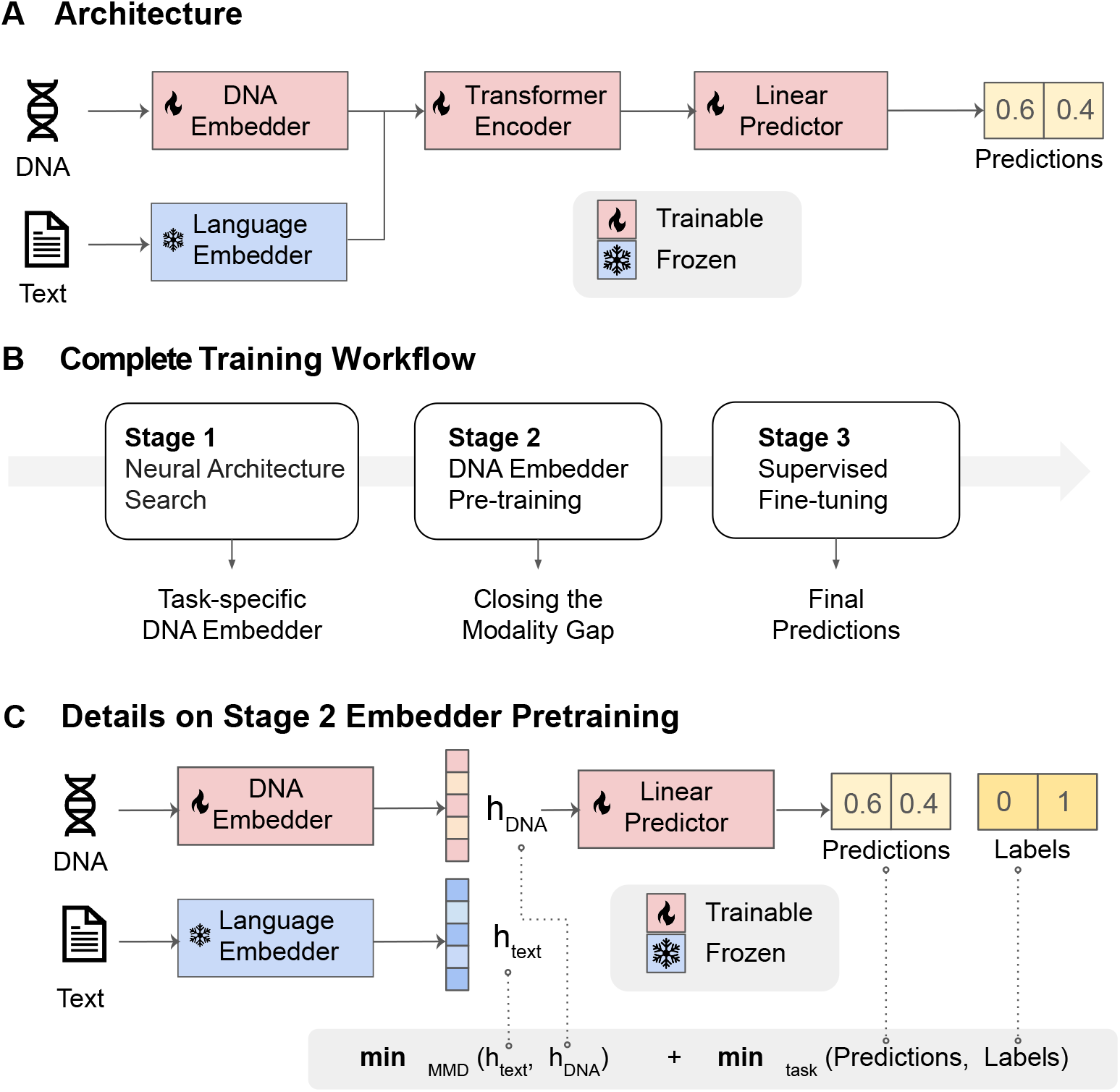
The overall workflow. **A**. The architecture of L2G. The architecture of the L2G model is composed of a CNN embedder, transformer layers from a pre-trained language model, and a linear predictor. **B**. The model is trained in three stages. In stage 1, L2G performs a Neural Architecture Search to optimize the embedder architecture for a given task. In stage 2, the CNN embedder is pre-trained to minimize the modality gap between DNA embeddings and language embeddings. In stage 3, the entire model is fine-tuned on task-specific data in a supervised manner by minimizing the task-specific loss between the final predictions and the true labels. **C**. In stage 2, L2G closes the modality gap by pertaining the embedder with a joint objective. The CNN embedder is trained with a joint objective, which simultaneously minimizes a) the distribution distance between language embeddings (*h*_text_) and DNA embeddings (*h*_DNA_), and b) the task-specific loss between predictions from Predictor 1 and the true labels.

The advantages of L2G are three-fold. First, by sidestepping pre-training, L2G is much more data and compute efficient. For a given downstream task, our three-stage workflow is comparable to vanilla fine-tuning in terms of required compute and data resources. All our experiments can be performed on a single A6000 GPU in a matter of hours by leveraging existing open-source language models, compared to days of training needed to develop genomic FMs from scratch. Second, L2G demonstrates better average performance than fine-tuned genomic FMs on various genomics benchmarks (**Fig**. 1B), including GenomicBenchmarks [13] and NucleotideTransformerBenchmarks [6]. Third, we show that L2G can tackle challenging regulatory activity prediction tasks, such as predicting developmental and house-keeping enhancer activity, where it consistently outperforms expert-designed models. Additionally, L2G learns relevant transcription factor motifs.

Overall, L2G leverages pre-trained LLMs for genomic prediction, achieving performance competitive with in-modal transfer on various genomic tasks while bypassing the massive costs associated with collecting and processing large amounts of unsupervised genomic sequencing data.

## Methods

### Cross-modal fine-tuning

Fine-tuning has been a highly effective technique for adapting pre-trained language models to various downstream tasks. However, most existing research focuses on *in-modal* adaptation, where the finetuning data originates from the same modality as the pre-training data but is tailored for a more specialized focus, such as sentiment analysis or text classification. In such cases, the model operates on the same type of input it was originally trained on. In contrast, *cross-modal* fine-tuning adapts a pre-trained model to work with data from an unseen modalities, such as using text-pre-trained LLMs to address biological questions. **Fig**. 1A illustrates the distinction between in-modal and cross-modal fine-tuning strategies for genomic tasks.

Cross-modal fine-tuning is more challenging than the in-modal fine-tuning due to the modality gap between the pre-training and target task data [14]. Bridge this gap often requires additional data alignment. For instance, to repurpose a pre-trained BERT model for predicting physicochemical and biomedical properties of protein sequences, Vinod et al. [10] introduced R2DL (Representation Reprogramming via Dictionary Learning), a token-level alignment method that learns a sparse linear mapping between English vocabulary embeddings and amino acid embeddings.

Recently, Shen et al [12] proposed a more general distributional alignment technique, ORCA, that adapts various pre-trained transformer models to diverse non-text, non-vision inputs. ORCA employs a convolutional neural network to transform input data into sequence features, minimizing the distribution distance between target data embeddings and standard English token embeddings prior to fine-tuning. ORCA achieves state-of-the-art results on three benchmarks containing over 60 datasets from 12 modalities, outperforming a wide range of general-purpose, automated machine learning (AutoML) and task-specific methods. However, genomics is one domain where ORCA does not show superior results [12]. Specifically, on DeepSEA – a well-known dataset for predicting functional effects of genomic sequence – ORCA falls behind non-pre-trained AutoML baselines. This motivates us to study how cross-modal alignment can be improved specifically for genomic prediction tasks.

Beyond ORCA, several other studies have also proposed different cross-modal fine-tuning strategies [15–20]. However, they are not tailored to biological domains. To address this gap, we aim to develop the first cross-modal fine-tuning framework specifically designed for genomic applications.

### Motivation for the framework

Since ORCA is the only previous method that has attempted genomics tasks, we thoroughly examine its workflow to identify limitations. Specifically, ORCA first creates custom embedder and predictor networks to support various tasks. The embedder is trained to minimize the optimal transport dataset distance (OTDD) between a target and proxy dataset, aiming to map the target dataset into the embedding space of the pre-trained model. Finally, the entire model – comprising the embedder, transformer, and predictor – is fully fine-tuned on the target task data to adapt the pre-trained model to the target modality.

However, ORCA has two major limitations when applied to genomics – both model-wise and training-wise. Model-wise, ORCA employs a universal CNN structure as the input embedder for all tasks. This embedder, consisting of a single-layer CNN with small kernel sizes and strides designed for computer vision, may not be well-suited for genomics tasks. It cannot effectively model the long-range dependencies of genomic sequences and may fail to capture important features from genomic datasets. To address this, we propose a redesigned embedder architecture tailored for genomics data.

Training-wise, by reproducing ORCA experiments on DeepSEA, we revealed that a lower alignment loss at the end of embedder training does not necessarily lead to better downstream performance on genomics tasks. This suggests a more complex dynamic between embedder training and fine-tuning than the ORCA paper indicated. For instance, in many cases, training the embedder for longer epochs can *hurt* the final performance on the target task. We hypothesize that this occurs because a single alignment loss is insufficient for effective embedder training; closer mapping to text embeddings can result in the loss of important class information in the target genomic dataset. To address this, we propose a new embedder training objective that jointly optimizes for both distribution alignment and downstream task performance.

By tailoring the embedder architecture and objective design to genomics data, we significantly improve empirical performance and develop a new cross-modal fine-tuning workflow, named L2G, to effectively adapt language models for genomic applications.

## Model design

### Problem Setup

A modality *M* consists of a feature space **𝒳**, a label space **𝒴**, and a joint probability distribution *P* (**𝒳**, **𝒴**). We focus on the cross-modal setting in this paper. That is, the target genomics modality *M*_*t*_ and source language modality *M*_*s*_ have different feature spaces, label spaces, and joint probability distributions, i.e., **𝒳** _*t*_ **𝒳** _*s*_, **𝒴**_*t*_*≠* **𝒴**_*s*_, and *P* (**𝒳** _*s*_, **𝒴**_*s*_)*≠ P* (**𝒳** _*t*_, **𝒴**_*t*_). Our goal is to adapt a model pre-trained in *M*_*s*_ to the tasks in *M*_*t*_.

Following previous work [12, 15], the model architecture of L2G is composed of three parts: a CNN embedder, a transformer encoder, and a linear predictor (**Fig**. 2A). The embedder maps input genomics data to an embedding space, the encoder extracts features from these embeddings, and the predictor maps the encoder output to the label space.

### Embedder

Denote *f* ^*s*^ as the source embedder of a language model, which transforms the source raw data **𝒳** _*s*_ into source language embeddings *h*^text^ = *R*^*N×D*^, where *N* denotes the embedding length and *D* denotes the embedding dimension. Following ORCA, we use the CoNLL-2003 dataset [21] as the reference dataset for text. This dataset contains nine classes for a named entity recognition (NER) task, from which we sampled 350 data points per class, resulting in a total of 3,150 data points. The source language embedder is taken from a pre-trained language transformer and remains frozen during the entire training process. Reference data is passed through the embedder to obtain reference embeddings for alignment.

Denote *f* ^*t*^ as the custom target embedder, which transforms the target genomics sequence data in **𝒳** ^*t*^ into target embeddings *h*^DNA^ = *R*^*N×D*^. The key to cross-modal transfer is to learn *f* ^*t*^ into map *h*^DNA^ into the shared representation space with *h*^text^. As mentioned above, existing work typically uses a generic small-kernel convolutional layer for *f* ^*t*^, which is unsuitable for modeling long-sequence genomics data. To address this, we propose using a larger, more capable dilated CNN as the backbone architecture for *f* ^*t*^. Previous studies have shown that dilated convolutions are effective for modeling DNA features in genomics tasks [22–25]. Unlike prior work, which fixes the convolution hyperparameters (e.g., kernel size and dilation rate) for the model architecture before seeing the task and data, we employ a data-driven approach that automatically learns the architecture configuration from the end task data (see later section). This new approach effectively improves downstream task performance.

### Transformer Encoder

The transformer encoder, denoted as as *g*, takes *h*^DNA^ as input and outputs intermediate representations (last hidden states) *h*^intermediate^ = *R*^*N×D*^. While L2G is compatible with various language models, we chose RoBERTa-base in our experiments in this work because its model size is smaller than or comparable to most transformer-based genomic FMs, such as DNABERT, DNABERT-2, Enformer, and Nucleotide Transformer-500M. This choice ensures a fair comparison of different methods. The embedding dimension *D* for RoBERTa-base is 768.

### Linear Predictor

The predictor, denoted as *p*^*t*^, takes *h*^intermediate^ as input and returns a task-specific output tensor. The goal of *p*^*t*^ is to map the learned representations to the desired output dimension. Following ORCA [12], we use average pooling along the sequence length dimension. A single linear layer then transforms the pooled outputs of the language models to produce the final prediction.

#### Model Training

L2G is trained in three stages: neural architecture search (NAS), embedder pre-training, and fine-tuning (see **Algorithm** 1 and **Fig**. 2B).

##### Algorithm 1 Pseudocode for the L2G workflow.

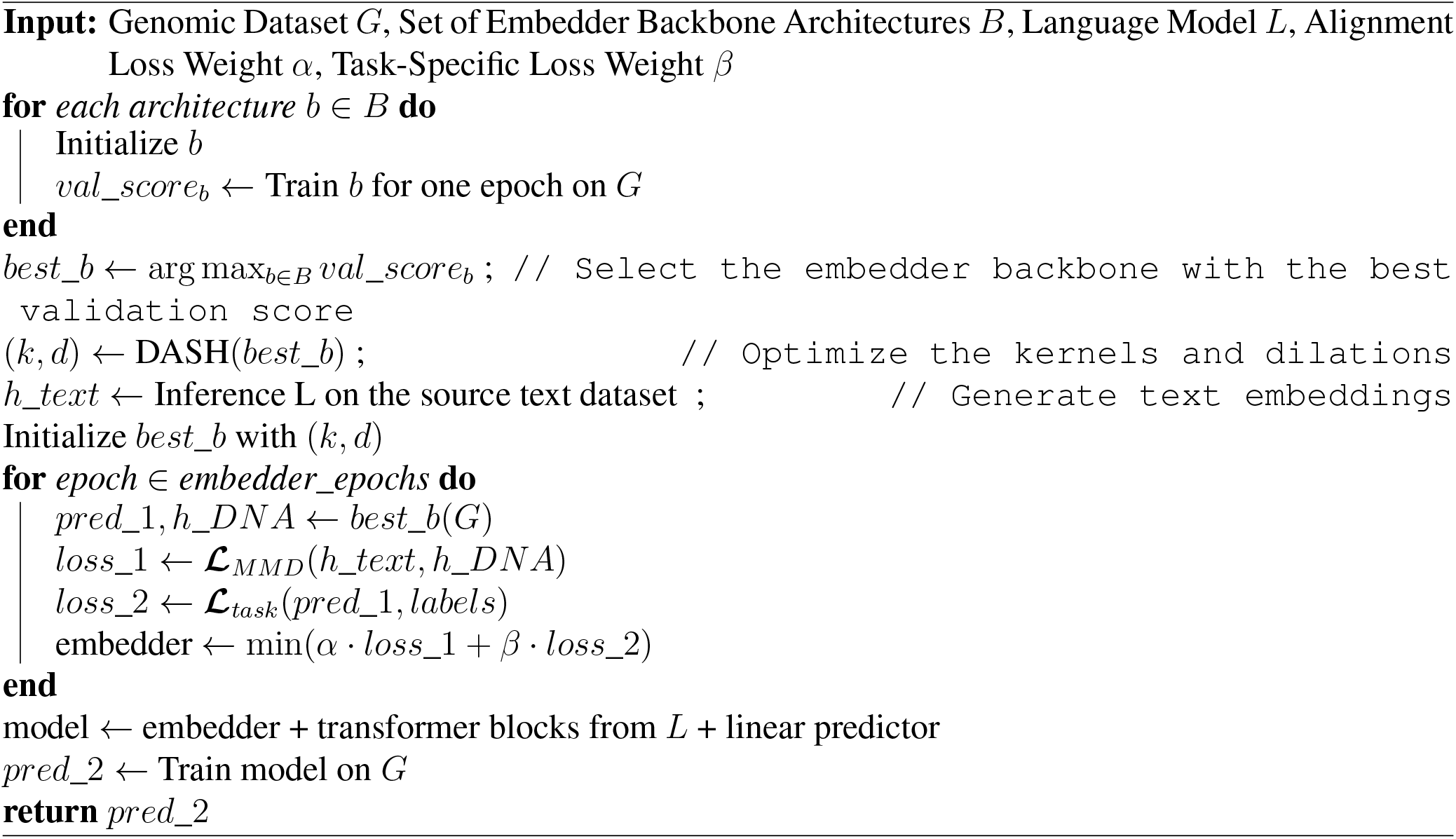

### Neural Architecture Search

Neural Architectural Search is a machine learning technique that automates the design of deep neural network architectures. Instead of relying on substantial manual efforts by human experts, NAS identifies architectures that perform well on a given task through algorithmic solutions [26]. Given its success in certain genomics applications [27], we utilize NAS to tailor embedder architectures to different tasks.

To achieve optimal performance across various downstream tasks, we use a two-step process for selecting the embedder network in L2G. First, we select an optimal *backbone* CNN architecture from a pre-defined search space. In this study, we consider ResNet [28] and UNet [29], both of which have been effectively applied in genomics. ResNet, with its deep residual connections, is well-suited for classification tasks that require capturing hierarchical features from sequential data. In contrast, UNet’s U-shaped encoder-decoder structure with skip connections makes it particularly effective for dense prediction tasks. Each architecture is trained for one epoch, and the one achieving the highest validation score is selected as the backbone.

Next, L2G applies NAS to optimize *layer operations* for the specific task. Specifically, we learn the optimal kernel size and dilation rate for each convolutional layer in the CNN using the DASH (Diverse-task Architecture Search) algorithm [30], which has demonstrated state-of-the-art performance among AutoML methods on the DeepSEA dataset. After this two-step process, both the backbone architecture and the convolutional layers of the embedder *f* ^*t*^ are tailored for the target task, effectively capturing meaningful target embeddings from genomics datasets.

### Embedder Pre-training

The embedder pre-training stage is critical for minimizing the modality gap between DNA and the pretrained language models, enabling cross-modal adaptation. We propose a joint objective to address the training limitations discussed earlier. The first objective minimizes the distribution distance between DNA and text data, performing modality alignment. Unlike ORCA [12], which uses OTDD loss, we utilize Maximum Mean Discrepancy (MMD) as the distance metric in L2G due to its better empirical performance in our ablation studies.

Additionally, we introduce a second objective – a task-specific loss (***ℒ***_task_) – during embedder pretraining. This enables the embedder to incorporate class information while performing distribution alignment. This task-specific loss is either cross-entropy loss for classication tasks or Mean Squared Error (MSE) for regression tasks. By including ***ℒ***_task_, the embedder learns to model the class information effectively when mapping genomics data to the language model’s embedding space. Training with only alignment loss but not task-specific loss can result in worse downstream performance, as shown in the ablation studies in the **Results** section.

In summary, the embedder pre-training objective is defined as:

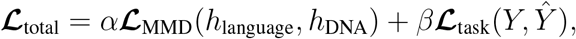

where *α* and *β* are weights for the MMD loss and task-specific loss, respectively. To optimize performance, we set up a scheduler for these weights, minimizing the task-specific loss first before minimizing the joint objective. We did this because empirically, introducing the alignment loss in later epochs of pre-training embedders achieves the best performance.

### Fine-tuning

After pre-training the embedder, the entire model – including embedder, transformer encoder, and linear predictor – is fine-tuned u sing t ask-specific lo ss on th e ta rget da ta. To op timize th e hyperparameter configuration (e.g., learning rate, dropout rate, weight decay) for fine-tuning, we use ASHA [31].

## Results

### Overview

Directly applying transformer models trained on natural language data to out-of-domain tasks like genomics can lead to the corruption of pre-trained weights, resulting in inefficiencies and inaccuracies due to the fundamental mismatch between the two modalities. To address this, we developed L2G, an effective and efficient workflow de signed to repurpose pr e-trained language mo dels fo r genomics tasks through cross-modal transfer learning. Unlike traditional in-modal transfer learning, where transformer models are first pre-trained on large-scale DNA sequencing data before fine-tuning, our approach is not only more competitive in prediction quality but also significantly more efficient. L2 G eliminates the need for large-scale self-supervised pre-training, reducing both data and computational requirements while still generalizing effectively across a variety of genomics tasks through fine-tuning.

We demonstrate the empirical effectiveness and efficiency of L2G through extensive experiments on two genomics benchmarks and a challenging regression task for enhancer activity prediction. Beyond presenting results on predictive accuracy, we assess L2G’s ability to learn relevant TF motifs and evaluate the efficacy of cross-modal fine-tuning through embedding analyses and ablation studies.

### L2G matches or outperforms fine-tuned genomic FMs

Predicting the regulatory function of non-coding DNA based on its sequence is crucial for prioritizing functional non-coding variants and remains a major challenge in genomics [32, 33]. We evaluated L2G on two existing benchmarks, Genomic Benchmarks [13] and the Nucleotide Transformer Benchmarks [6], to demonstrate its generalizability and efficacy.

We first evaluated L2G on the Nucleotide Transformer Benchmarks [6], one of the most widely used benchmarks for genomic FMs. This benchmark suite includes eighteen tasks for predicting regulatory elements from four categories: enhancers, promoters, epigenetic marks, and splice sites from DNA sequences with lengths ranging from 300 to 600 bp (**Table** S3). We compared L2G against several representative genomic FMs, including Enformer [34], DNABERT-1 [5], DNABERT-2 [8], HyenaDNA (1kb) [7], Nucleotide Transformer - Multi-species (2.5B) and Caduceus-ph [35], which outperforms Caduceus-ps in most of the tasks [6]. All these models have been pre-trained and then finetuned.

The complete benchmarking results are displayed in **Table** S4 and **Fig**. 3. L2G achieves the best results on ten tasks and ranks second on six others. It demonstrates a clear advantage in predicting histone marks and enhancers from DNA sequences, outperforming all other genomic FMs. For promoter and splice site prediction tasks, the Nucleotide Transformer is the top-performing model, followed by L2G. It is worth noting that the Nucleotide Transformer is the largest model evaluated on this benchmark, with 2.5 billion parameters and the most extensive pre-training data, while L2G uses significantly fewer parameters and no pre-training data. Overall, L2G achieves the highest average test score on the Nucleotide Transformer benchmark with the least training (pre-training + fine-tuning) data, underscoring its ability to generalize effectively across tasks even with limited data.

**Figure 3:**
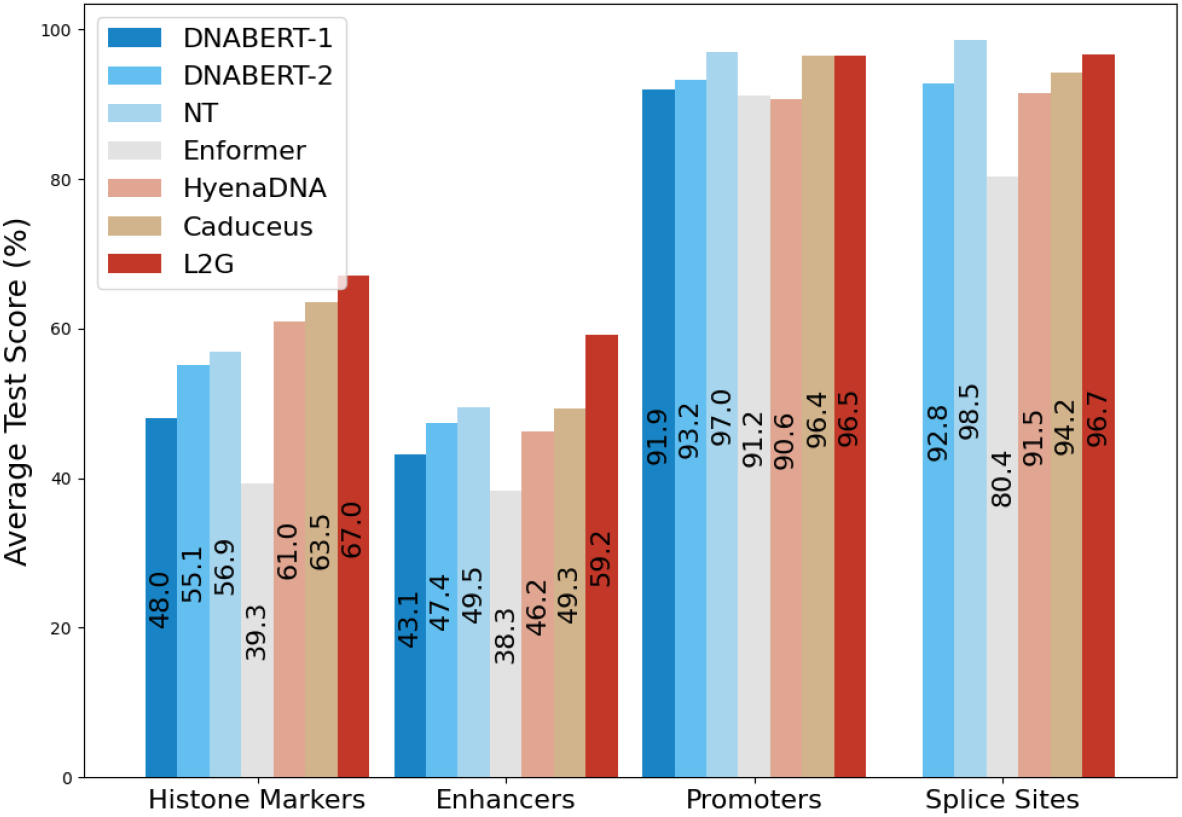
Average performance across task types (histone marks, enhancers, promoters, and splice sites) on the Nucleotide Transformer benchmarks. Each bar represents the average test score of a model. L2G outperforms all other models on histone marks and enhancers tasks and ranks second on promoters and splice sites prediction tasks. Notably, the bar for DNABERT-1 in the splice sites category is missing because it could not be trained on two splice site prediction tasks.

The Genomic Benchmarks dataset [13] includes eight classification tasks: seven binary and one three-way classification task. These tasks focus on predicting regulatory elements such as promoters, enhancers, and open chromatin regions from several species, including humans and *Drosophila* [7, 13, 36]. The inputs sequences have median lengths ranging from 200 to 2,381 bp. This benchmark includes three baselines: a supervised CNN model, a supervised transformer model, and a fine-tuned genomic FM, HyenaDNA [7].

The results are shown in **Table** S5. L2G outperforms all other models in five out of eight tasks and is the second best in the remaining three, slightly behind HyenaDNA. As demonstrated by the aggregated results using performance profiles **Fig**. S1C, L2G achieves the best overall performance on the Genomic Benchmarks.

Overall, L2G matches or outperforms fine-tuned genomic FMs across a variety of regulatory element prediction tasks. This is particularly significant as L2G does not rely on extensive pre-training on unsupervised DNA data, a standard practice for most genomic FMs. These results highlight the efficacy of the cross-modal transfer learning approach employed by L2G, which effectively leverages the pre-trained knowledge embedded in language models to address genomics tasks.

### L2G reveals transcriptional factor motifs

While we have demonstrated the strong benchmark performance of L2G, we also sought to showcase its utility in downstream applications, such as discovering functional regulatory syntax. Here, we focused on a regression task to predict the activities of developmental and housekeeping enhancers [37] from DNA sequences. Using the DeepSTARR dataset [37], which predicts enhancer activity for two distinct promoters in *Drosophila* S2 cells, we compared L2G to baseline methods.

The comparison included the DeepSTARR model, an adaptation of the Basset convolutional neural network [38], as well as several genomic FMs. Full results are presented in **Table** S6. **Fig**. 4A shows that predictions by L2G align well with measured values for both developmental (PCC=0.66) and housekeeping (PCC=0.76) enhancers. Compared to other models, L2G outperforms all others in the housekeeping enhancer prediction task and ranks second in the developmental enhancer prediction task, slightly behind DeepSTARR (PCC=0.68).

**Figure 4:**
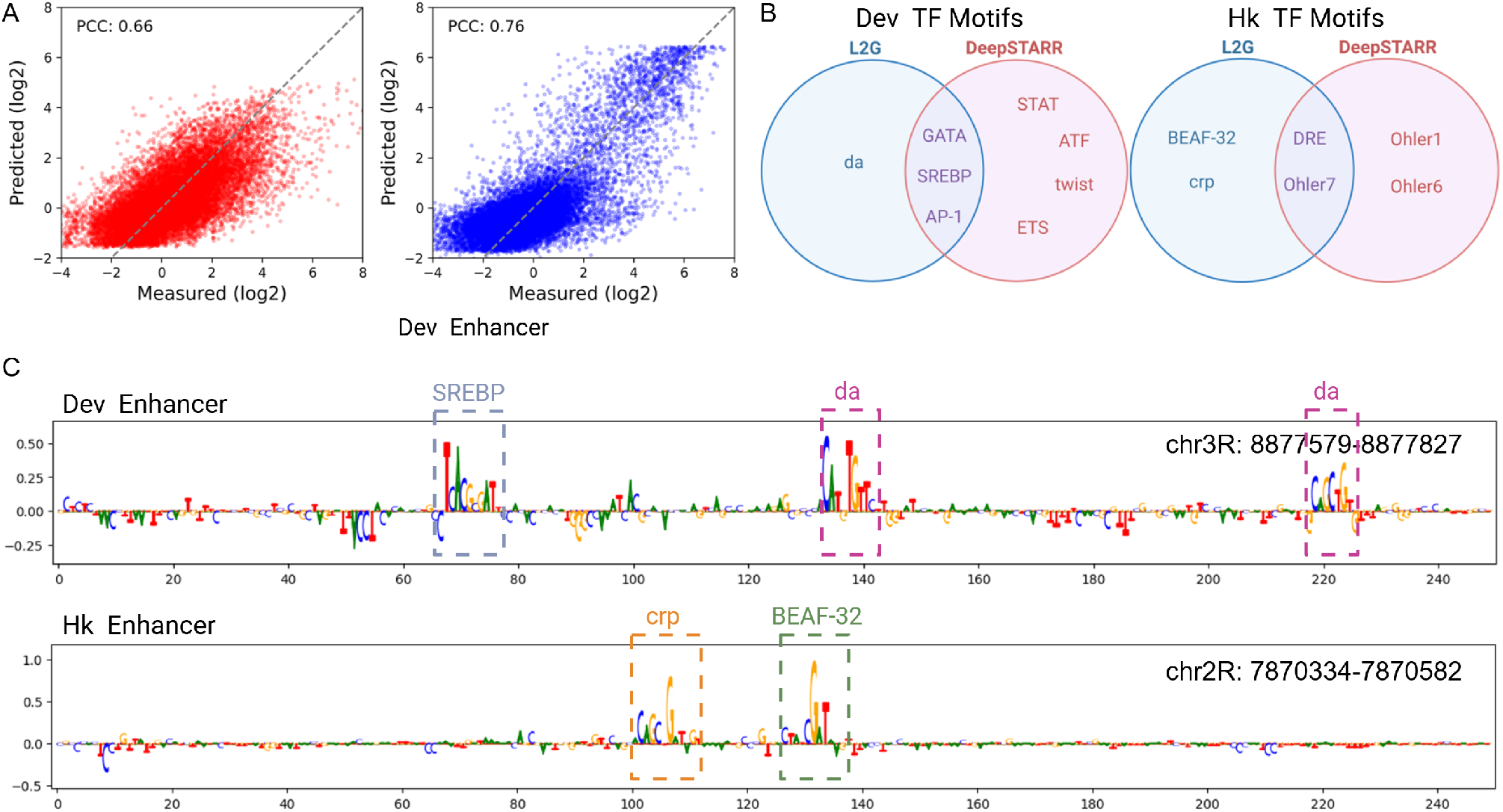
L2G predicts enhancer activities and reveals TF motifs. **A**. Performance of L2G on predicting developmental and housekeeping enhancer activity from DNA sequences in *Drosophila* S2 cells, measured using Pearson Correlation Coefficient (PCC). Scatter plots show predicted vs. observed enhancer activity for developmental (left) and housekeeping (right) enhancers. **B**. Venn diagrams indicating the common and unique TF motifs identified by L2G (blue) and DeepSTARR (red) for developmental (left) and housekeeping enhancers (right). Motif identified by DeepSTARR were retrieved from [37], with unknown and redundant motifs excluded. While both methods achieve high PCC values in predicting enhancer activities, they identify different sets of motifs. **C**. Nucleotide contribution scores for strong developmental (top) and housekeeping (bottom) enhancer sequences, respectively.

To determine whether L2G learned regulatory syntax, we quantified how each nucleotide contributes to predicted enhancer activities using DeepLiftShap [39] and identified predictive sequence patterns with TF-Modisco-lite [40]. Full motif results are shown in **Figs**. S3 and **Figs**. S4. Interestingly, although both L2G and DeepSTARR achieve high PCC values in predicting developmental and housekeeping enhancer activities, they identified different sets of TF motifs (**Fig**. 4B).

For developmental TF motifs, both models identified AP-1, GATA and SREBP, but L2G uniquely revealed the da motif. The daughterless (da) gene, part of the basic helix-loop-helix (bHLH) family, is essential for several developmental pathways, including sex determination and neurogenesis [41]. For housekeeping TF motifs, L2G uniquely identified BEAF-32 and CRP. The *Drosophila* Boundary Element-Associated Factor (BEAF) of 32kDa primarily binds near the promoters of numerous house-keeping genes, contributing to chromatin domain boundary activity and promoter function [42, 43]. Similarly, the CRP motif may be associated with housekeeping promoters [44].

We acknowledge that differences in identified motifs may arise from several factors. First, we used different implementations of DeepLIFTShap and TF-MoDISco algorithms, as DeepSTARR is based on Keras and TensorFlow, while our implementation uses PyTorch. These framework differences can contribute to variations in motif interpretation and sensitivity. Additionally, DeepSTARR’s simpler CNN-based architecture is inherently easier to interpret than L2G’s CNN-transformer hybrid model, and hyperparameter variations could also affect motif detection.

Nevertheless, we demonstrate that L2G can predict enhancer activities and reveal relevant TF motifs associated with developmental and housekeeping enhancers, suggesting it is effective in identifying important sequence patterns for prediction tasks.

### L2G closes the modality gap

L2G bridges the modality gap through joint loss optimization during the embedder pre-training step, simultaneously aligning distributions between text and DNA while optimizing downstream task performance. This approach addresses the challenge where directly fine-tuning language models on genomic tasks often results in weight shifts and poor performance. To evaluate the effectiveness of this strategy, we examined the learned representations of the target modality data generated by different fine-tuning approaches. Specifically, we selected three binary classification tasks from the Nucleotide Transformer benchmark and visualized the learned embeddings for the two classes. We also calculated the Silhouette Score, which quantifies cluster separation. Scores range from -1 to +1, with +1 indicating well-separated clusters, 0 suggesting overlapping clusters, and -1 indicating incorrect class assignments [45]. Across all three tasks, L2G achieved higher positive Silhouette Scores (**Fig**. S2), demonstrating improved class separation compared to vanilla fine-tuning. This clear distinction in embedding space resulted in better performance.

To better understand the factors contributing to the success of cross-modal fine-tuning in L2G, we conducted three ablation studies on selected tasks from the Nucleotide Transformer benchmark, H3, enhancer, and promoter_tata. The full empirical results are provided in **Tables** S7, S8, and S9.

First, to validate the importance of using a pre-trained LLM backbone, we compared two configurations: using a pre-trained RoBERTa-base transformer vs. a randomly initialized backbone. Next, we examined the impact of loss functions during the embedder pre-training step by comparing joint loss optimization with task-specific loss alone and MMD loss alone. Finally, we evaluated the optimal choice of embedder architecture by comparing the neural architecture search method DASH, used in L2G, with two alternatives: the domain-specific CNN model DeepSEA and the embedder from the original ORCA model [12].

The results of these ablation studies showed that the pre-trained transformer, joint loss optimization, and the DASH-based embedder consistently outperformed their respective alternatives (**Fig**. 5). These findings highlight the critical role of pre-training, optimized loss functions, and robust embedder architecture in the success of L2G.

**Figure 5:**
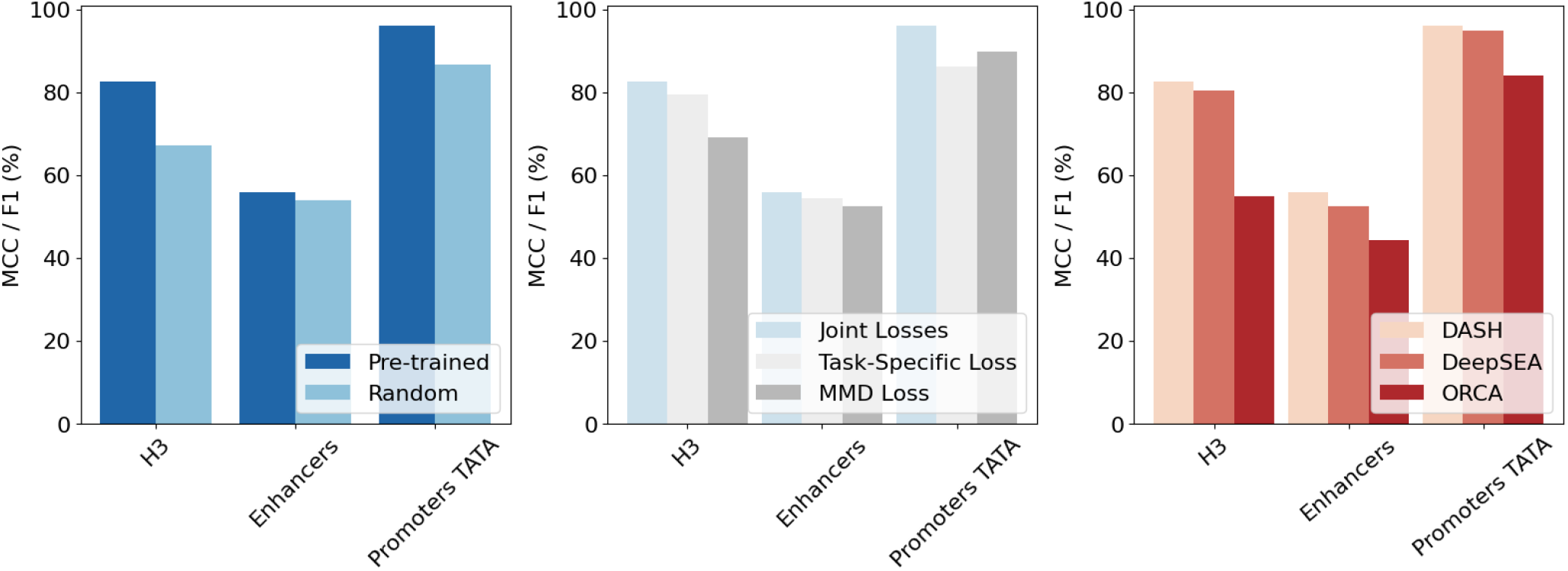
Ablation results for three selected tasks in the Nucleotide Transformer benchmark. Left: Performance comparison between a pre-trained RoBERTa-base transformer and a randomly initialized backbone during pretraining. Middle: Middle: Evaluation of different loss functions during embedder pretraining, comparing joint loss optimization, task-specific loss alone, and MMD loss alone. Right: Comparison of embedder architectures, including DASH, DeepSEA, and the original ORCA embedder.

## Discussion

In this work, we investigate the efficacy of cross-modal transfer in genomics. By analyzing a general-purpose cross-modal fine-tuning method, we identified key limitations in both architectural design and objective function. To address these challenges, we introduced L2G, a new method that incorporates a carefully designed architecture and improved alignment between different modalities. This enables L2G to harness the capabilities of pre-trained language models for genomics tasks. Our evaluations across multiple genomics tasks demonstrate superior average performance compared to fine-tuned genomic FMs and domain-specific expert models, notably without requiring large-scale pre-training.

The success of our cross-modal approaches raises important questions: Is the current pre-training approach in genomics the most effective? Do we truly need vast amounts of unsupervised genomics data for pre-training? By leveraging pre-trained language models, L2G bypasses the need for extensive unsupervised pre-training, reducing computational and data demands while still achieving competitive performance with in-modal transfer. This challenges the conventional approach of building domain-specific FMs from scratch, suggesting that language models originally developed for NLP can be repurposed for seemingly unrelated domains like genomics. This opens new avenues for cross-disciplinary applications of LLMs, highlighting their versatility and questioning the necessity of developing entirely new models for every domain.

A few contemporaneous studies have also raised concerns about the effectiveness of current genomic FMs. For example, one study observed that genomic FMs offer little to no advantage over traditional models based on one-hot encoded sequences [46]. Another work found that a supervised-only pipeline named DASHA surpassed the latest genomic FMs on the Nucleotide Transformer benchmark [47]. Notably, L2G outperforms DASHA on 11/18 tasks in the benchmark and reaches a better average score. At the time of this writing, L2G is the only fine-tuning based approach that outperforms such strong supervised baselines on the Nucleotide Transformer benchmark, despite not being pretrained on genomics data. Our work along with these contemporaneous studies collectively challenge the prevailing pretraining-then-fine-tuning paradigm for genomic FMs and highlight the need to rethink their development and applications.

Our study has several limitations. First, our evaluations did not cover a broad range of genomics tasks, including long-range prediction tasks that involve more complex interactions and regulatory mechanisms. Future work could extend L2G’s application to more complex genomic tasks, such as sequencebased gene expression prediction [22, 34]. Second, we have not explored whether the scaling laws common in NLP apply to cross-modal transfer learning for genomics. It remains to be seen whether using increasingly larger language models (trained on natural language) would yield proportionally better performance. Thirdly, interpretability remains a challenge [48]. While we have used ablation studies and embedding analyses to explain L2G’s effectiveness, the underlying mechanisms and interpretability of L2G require further investigation. Lastly, our current cross-modal transfer approach relies on finetuning. Future work could explore combining cross-modal transfer with continued pre-training [49], leveraging both unsupervised text and genomic data to potentially further enhance the performance of domain-specific FMs, albeit at the cost of increased data and computational requirements.

In summary, L2G demonstrates the potential of cross-modal transfer learning to address genomics tasks effectively and efficiently, providing a compelling case for leveraging existing pre-trained models from natural language rather than building domain-specific ones from scratch. This work lays the foundation for further advancements in cross-disciplinary applications of pre-trained language models, extending their utility to a diverse range of biological problems.

## Code Availability

The source code of L2G can be accessed at: https://github.com/wenduocheng/L2G.

## Data Availability

In this work, we utilized several public datasets.

- The Genomic Benchmark is available at: https://github.com/ML-Bioinfo-CEITEC/genomic_benchmarks.
- The Nucleotide Transformer benchmarks can be downloaded from HuggingFace at: https://huggingface.co/datasets/InstaDeepAI/nucleotide_transformer_downstream_tasks.
- The DeepSTARR dataset is available on Zenodo at: https://doi.org/10.5281/zenodo.5502060.

## Acknowledgments

This work was supported in part by the National Institutes of Health Common Fund 4D Nucleome Program grant UM1HG011593 (J.M.), National Institutes of Health Common Fund Cellular Senescence Network Program grant UH3CA268202 (J.M.), National Institutes of Health grants R01HG007352 (J.M.), R01HG012303 (J.M.), R21DA061481 (J.M.), and U24HG012070 (J.M.), as well as National Science Foundation grants IIS1705121 (A.T.), IIS1838017 (A.T.), IIS2046613 (A.T.), and IIS2112471 (A.T.). J.M. was additionally supported by a Guggenheim Fellowship from the John Simon Guggenheim Memorial Foundation, a Google Research Award, and a Single-Cell Biology Data Insights award from the Chan Zuckerberg Initiative. A.T. received additional funding from Meta, Morgan Stanley, Amazon, Google, and Scribe. Any opinions, findings, conclusions, or recommendations expressed in this material are those of the author(s) and do not necessarily reflect the views of any of these funding agencies.

## Author Contributions

Conceptualization: W.C., J.S., A.T., and J.M.; Code: W.C. and J.S.; Investigation: W.C., J.S., M.K., A.T., and J.M.; Writing: W.C., J.S., M.K., A.T., and J.M.; Funding Acquisition: J.M and A.T.

