## Supplemental Information for "L2G: Repurposing Language Models for Genomics Tasks"

#### Supplementary Notes

##### Pre-training resources of various DNA FMs

**Table** S1 provides a comparative summary of the computational resources, model parameters, and pre-training data with various DNA FMs.

##### Description of the downstream tasks

Genomic Benchmarks dataset consists of eight classification tasks: seven binary and one three-way, focusing on regulatory elements such as promoters, enhancers, and open chromatin regions from several species, including humans, mouse (*Mus musculus*), and roundworm (*C. elegans*) [1]. A three-layer CNN serves as the baseline model in this benchmark, and the HyneaDNA study [2] included a supervised trained transformer baseline. Inputs are DNA sequences with lengths between 200 to 500 bp, except for the Mouse Enhancer Ensembl dataset, which has the longest inputs (median 2,381 bp; maximum 4,707 bp). Metadata for the tasks included in the Genomic Benchmarks [1] is provided in **Table S2**.

Nucleotide Transformer Benchmarks dataset is another widely used benchmark for evaluating DNA FMs. This benchmark suite, introduced alongside the Nucleotide Transformer [3], evaluates DNA FMs on 18 classification tasks such as predicting regulatory elements for enhancers (human), promoters (human/mouse), epigenetic marks (yeast), and splice sites (human/multispecies) from DNA sequences 300-600 bp long. Performance metrics for several models – including Enformer [4], DNABERT-1 [5], DNABERT-2 [6], HyenaDNA [7], and Nucleotide Transformer [3] – are included, along side results the recent Caduceus-Ph [8]. Metadata for the tasks included in the Nucleotide Transformer Benchmarks [3] is presented in **Table S3**.

Developmental and Housekeeping Enhancer Activity Predictions is a two-class regression task that predicts enhancer activities for housekeeping and developmental enhancers in *Drosophila* S2 cells using 249 bp sequences. The dataset, sourced from the DeepSTARR project [9], includes a CNN model baseline. The evaluation metric is the Pearson Correlation Coefficient (PCC).

##### Complete results

The complete results on the Nucleotide Transformer Benchmarks are shown in **Table S4** and **Fig. S1A**. We used a batch size of 64 and cross-entropy loss across all datasets. Test scores for other DNA FMs are from Supplementary Table 6 of Dalla-Torre et al.[3]. Results for Caduceus-Ph [8], which outperforms the Caduceus-Ps on 17 out of 18 tasks, are included. To provide a holistic comparison of methods across all datasets, we utilized performance profiles [10]. Each curve shows the proportion of problems it solves within varying thresholds of a performance factor $\tau$. As shown in **Fig. S1B**, L2G achieves the best or second-best performance across all tasks.

The complete results on the Genomic Benchmarks are shown in **Table S5**. We used a batch size of 64 and cross-entropy loss across all datasets. We trained the CNN and HyenaDNA (32k) baselines, while results for the transformer baseline were obtained from the HyenaDNA paper [2], as the code is not open-sourced. We also computed the performance profiles for the results on the Genomic Benchmarks. As shown in **Fig. S1C**, L2G achieves top performance across all tasks.

The complete results for the Developmental and Housekeeping Enhancer Activity Prediction Task are presented in **Table S6**. For this task, we used a batch size of 128 and mean squared error (MSE) loss. To benchmark the performance of L2G, we included the expert model, DeepSTARR, which was a CNN specifically designed for this task. Additionally, we compared with two DNA FMs, HyenaDNA (32k) and Nucleotide Transformer (v2, 500M). PCC was used as the evaluation metric.

##### Embedding analysis

To evaluate the quality of the learned representations of the target modality data, we analyzed the embeddings generated by L2G and compared them to those obtained through vanilla fine-tuning. This analysis was conducted on three binary classification tasks from the Nucleotide Transformer benchmarks: H3, enhancers, and promoter_tata. We visualized the embeddings of the two classes in each task using t-SNE, as shown in **Fig. S2**. This visualization provided a qualitative assessment of how well the embeddings separate the classes in a reduced-dimensional space. To quantitatively measure cluster separation, we calculated the Silhouette Score for each set of embeddings. The Silhouette Score ranges from -1 to +1, where +1 indicates well-separated clusters, 0 signifies overlapping clusters, and -1 suggests incorrect class assignments [11]. Across all three datasets, L2G consistently achieved higher Silhouette Scores compared to vanilla fine-tuning, demonstrating its superior ability to produce distinct class separations in the embedding space.

##### Ablation studies

We conducted a series of ablation studies to evaluate the contributions of different components in our method. Specifically, we assessed the impact of pre-trained transformers, the losses during embedder pretraining, and the embedder architecture. The results on three tasks from the Nucleotide Transformer benchmarks are shown. Detailed results are provided in **Table** S7 for ablation study for the pre-training Body, **Table** S8 for ablation study for the losses during embedder pre-training, and **Table** S9 for the embedder architecture.

##### Motif analysis

We calculated the nucleotide contribution scores using a DeepLiftShap method from Catum GitHub Repository for developmental and housekeeping enhancer activities. DeepLiftShap combines the DeepLIFT [12] algorithm with SHAP (SHapley Additive exPlanations) values to attribute model predictions to input features by calculating the contribution relative to a reference baseline [13]. Each feature is a nucleotide at a specific position.

Following the DeepSTARR methodology [9], we used 100 dinucleotide-shuffled versions of each input sequence as baseline sequences. Hypothetical importance scores for each sequence were multiplied by its one-hot encoded matrix to derive the final nucleotide contribution scores.

Motifs were identified using TF-Modisco-lite, a more efficient implementation of TF-Modisco [14], on the nucleotide contribution scores for each enhancer type separately [9]. For motif annotation, we downloaded two reference databases for *Drosophila*: OnTheFly [15] and FlyFactorSurvey [16], from the MEME suite [17], and compared motifs using TOMTOM [18]. Motifs with fewer than 30 seqlets were discarded. The resulting motifs for developmental and housekeeping enhancers are visualized in **Fig.** S3 and **Fig.** S4, respectively,

##### Implementation details

For all tested datasets, we applied data during training by randomly shifting input sequence by up to 3 bp and reverse-complementing sequences. During testing, predictions from the forward and reverse complement sequences were averaged. This approach is commonly used in genomics to improve the prediction accuracy of deep learning models [4, 19].

### Supplementary Tables and Figures

| Model | Params | GPUs | Wall clock | Pre-training Data |
|---|---|---|---|---|
| DNABERT [5] | 110M | 8-GTX 2080ti-11GB | 25 days | 3.2B |
| DNABERT-2 [6] | 117M | 8-GTX 2080ti-11GB | 14 days | 32.5B |
| Enformer [4] | 252M | 64-TPU v3 cores-32TB | 3 days | 14.1B |
| Nucleotide Transformer [3] | 2.5B | 128-A100-80GB | 28 days | 174B |
| HyenaDNA [2] | 32K | 1-A100-40GB | 80 mins | 3.2B |
| Caduceus [8] | 1.9M | ? | ? | 35B |

**Table S1:** Pre-training resources and data of various DNA foundation models. The pre-training data is reported in nucleotides. The computing resources required for pre-training Caduceus are unknown. Note that Enformer is a supervised model trained for the gene expression prediction task and is not pre-trained on unsupervised genomic sequencing data. However, we included it here as it was used as a DNA FM baseline in the Nucleotide Transformer benchmark.

| Dataset | Samples | Classes | Max Length | Metric |
|---|---|---|---|---|
| dummy_mouse_enhancers_ensembl | 1,210 | 2 | 4,707 | Accuracy |
| demo_coding_vs_intergenomic_seqs | 100,000 | 2 | 200 | Accuracy |
| demo_human_or_worm | 100,000 | 2 | 200 | Accuracy |
| human_enhancers_cohn | 27,791 | 2 | 500 | Accuracy |
| human_enhancers_ensembl | 154,842 | 2 | 573 | Accuracy |
| human_ensembl_regulatory | 289,061 | 3 | 802 | Accuracy |
| human_nontata_promoters | 36,131 | 2 | 251 | Accuracy |
| human_ocr_ensembl | 174,756 | 2 | 593 | Accuracy |

**Table S2:** Description of datasets in Genomic Benchmarks. Each dataset is described by the name, the total number of samples, the number of target classes, the maximum sequence length in nucleotides, and the evaluation metric used.

| Dataset | Samples | Classes | Max Length | Metric |
|---|---|---|---|---|
| H3 | 13,468 | 2 | 500 | MCC |
| H3K4me1 | 28,509 | 2 | 500 | MCC |
| H3K4me2 | 27,614 | 2 | 500 | MCC |
| H3K4me3 | 33,119 | 2 | 500 | MCC |
| H3K9ac | 25,003 | 2 | 500 | MCC |
| H3K14ac | 29,743 | 2 | 500 | MCC |
| H3K36me3 | 31,392 | 2 | 500 | MCC |
| H3K79me3 | 25,953 | 2 | 500 | MCC |
| H4 | 13,140 | 2 | 500 | MCC |
| H4ac | 30,685 | 2 | 500 | MCC |
| enhancer | 14,968 | 2 | 200 | MCC |
| enhancer_types | 14,968 | 3 | 200 | MCC |
| promoter_all | 53,276 | 2 | 300 | F1 |
| promoter_tata | 5,517 | 2 | 300 | F1 |
| promoter_non_tata | 47,759 | 2 | 300 | F1 |
| Splice sites all | 27,000 | 2 | 400 | Accuracy |
| splice_sites_acceptor | 19,961 | 2 | 600 | F1 |
| splice_sites_donor | 19,775 | 2 | 600 | F1 |

**Table S3:** Description of datasets in Nucleotide Transformer Benchmarks. Each dataset is described by the name, the total number of samples, the number of target classes, the maximum sequence length in nucleotides, and the evaluation metric used. Metrics include MCC (Matthews Correlation Coefficient), F1 score, and accuracy, as used in the Nucleotide Transformer study [3].

| Dataset | NT | Enformer | DNABERT-1 | DNABERT-2 | HyenaDNA | Caduceus-Ph | L2G |
|---|---|---|---|---|---|---|---|
| *Histone Markers* | | | | | | | |
| H3 | 79.3 | 72.4 | 76.3 | 78.5 | 78.1 | <u>81.5</u> | **82.5** |
| H3K4me1 | <u>54.1</u> | 29.1 | 39.6 | 51.2 | 51.2 | 52.3 | **58.6** |
| H3K4me2 | 32.4 | 20.7 | 28.2 | 33.3 | 45.5 | <u>48.7</u> | **56.2** |
| H3K4me3 | 40.8 | 15.6 | 25.8 | 35.3 | <u>55.0</u> | 54.4 | **66.3** |
| H3K9ac | 54.7 | 41.5 | 50.5 | 54.5 | 58.6 | <u>62.2</u> | **65.1** |
| H3K14ac | 53.8 | 28.4 | 40.3 | 51.5 | 60.8 | <u>63.1</u> | **69.4** |
| H3K36me3 | <u>61.8</u> | 34.5 | 47.4 | 59.1 | 61.4 | 60.1 | **68.8** |
| H3K79me3 | 62.3 | 49.8 | 57.8 | 61.5 | 66.9 | <u>69.7</u> | **70.7** |
| H4 | <u>80.8</u> | 73.5 | 78.4 | 79.7 | 76.3 | **81.1** | 78.8 |
| H4ac | 49.2 | 27.5 | 35.9 | 46.5 | 56.4 | <u>62.1</u> | **65.1** |
| **Average** | 56.9 | 39.3 | 48.0 | 55.1 | 61.0 | <u>63.5</u> | **67.0** |
| *Enhancer* | | | | | | | |
| Enhancers | <u>54.5</u> | 45.4 | 49.5 | 52.5 | 52.0 | 54.6 | **55.8** |
| Enhancer types | <u>44.4</u> | 31.2 | 36.7 | 42.3 | 40.3 | 43.9 | **62.6** |
| **Average** | <u>49.5</u> | 38.3 | 43.1 | <u>47.4</u> | 46.2 | 49.3 | **59.2** |
| *Promoter* | | | | | | | |
| Promoter TATA | **95.9** | 91.8 | 91.0 | 90.9 | 87.9 | 95.3 | <u>96.0</u> |
| Promoter non-TATA | **97.7** | 90.9 | 92.4 | 94.3 | 91.9 | 96.9 | <u>97.2</u> |
| Promoter all | **97.5** | 90.9 | 92.2 | 94.3 | 91.9 | <u>97.0</u> | 96.2 |
| **Average** | **97.0** | 91.2 | 91.9 | 93.2 | 90.6 | 96.4 | <u>96.5</u> |
| *Splice Sites* | | | | | | | |
| Splice sites all | **98.2** | 77.2 | 96.2 | 90.9 | 93.4 | 94.0 | <u>97.9</u> |
| Splice sites acceptors | **98.6** | 82.9 | / | 94.9 | 91.6 | 93.7 | <u>96.4</u> |
| Splice sites donors | **98.7** | 81.2 | / | 92.5 | 89.4 | 94.8 | <u>95.5</u> |
| **Average** | **98.5** | 80.4 | / | 92.8 | 91.5 | 94.2 | <u>96.7</u> |

**Table S4:** The performance of each model on the Nucleotide Transformer Benchmarks. Metrics used by task: MCC for histone markers, F1-score for enhancers and splice site acceptors/donors, and accuracy for splice site all. Bold indicates the best performance, and underline indicates the second-best. NT stands for nucleotide transformer (multispecies, 2.5B version). The results for other baselines are retrieved from the Nucleotide Transformer paper [3]. DNABERT-1 could not be trained on two splice site prediction tasks because the input sequence length exceeded the maximum context length allowed by DNABERT-1.

| Dataset | CNN | Transformer | HyenaDNA | L2G |
|---|---|---|---|---|
| Mouse Enhancers | 72.8 | 80.1 | **82.6** | <u>79.3</u> |
| Coding vs Intergenomic seqs | 88.2 | 88.8 | **89.6** | **91.7** |
| Human vs Worm | 92.8 | 95.6 | <u>96.5</u> | **96.5** |
| Human Enhancers Cohn | 71.6 | 70.5 | <u>73.0</u> | **73.2** |
| Human Enhancers Ensembl | 80.2 | 83.5 | <u>86.9</u> | **88.4** |
| Human Ensembl Regulatory | 94.0 | 91.5 | 92.0 | **93.9** |
| Human Nontata Promoters | 85.8 | 87.7 | **94.3** | <u>92.9</u> |
| Human OCR Ensembl | 67.8 | 73.0 | **79.1** | <u>78.9</u> |
| **Average** | 81.7 | 83.8 | <u>86.8</u> | **86.9** |

**Table S5:** The performance of each model on the Genomic Benchmark dataset. The evaluation metric is accuracy (the higher, the better). Bold indicates the best performance, and underline indicates the second-best.

| Dataset | HyenaDNA | Nucleotide Transformer | DeepSTARR | L2G |
|---|---|---|---|---|
| dev | 0.57 | 0.64 | **0.68** | 0.66 |
| hk | 0.65 | 0.75 | 0.74 | **0.76** |
| Mean | 0.61 | 0.70 | **0.71** | **0.71** |

**Table S6:** The performance of each model on the *Drosophila* enhancers prediction regarding the developmental (dev) and housekeeping activity (hk).

| Dataset | Pre-trained | Random |
|---|---|---|
| H3 | 82.5 | 67.1 |
| Enhancers | 55.8 | 54.0 |
| Promoters TATA | 96.0 | 86.7 |

**Table S7:** Performance comparison between pre-trained RoBERTa and randomly initialized RoBERTa on three selected datasets on the Nucleotide Transformer benchmark. The results are reported as MCC (higher is better).

| Dataset | Joint losses | Task-specific loss only (alpha=0) | MMD loss only (beta=0) |
|---|---|---|---|
| H3 | 82.5 | 79.5 | 69.0 |
| Enhancers | 55.8 | 54.5 | 52.4 |
| Promoters TATA | 96.0 | 86.2 | 89.8 |

**Table S8:** Performance comparison of different losses used during the embedder pre-training stage on three selected datasets on the Nucleotide Transformer benchmark.

| Dataset | DASH | DeepSEA | ORCA |
|---|---|---|---|
| H3 | 82.5 | 80.5 | 54.9 |
| Enhancers | 55.8 | 52.4 | 44.3 |
| Promoters TATA | 96.0 | 94.9 | 84.0 |

**Table S9:** Performance comparison between the embedder architecture of DASH, DeepSEA (unsearched), and ORCA (conv1d) on three selected datasets on the Nucleotide Transformer benchmark.

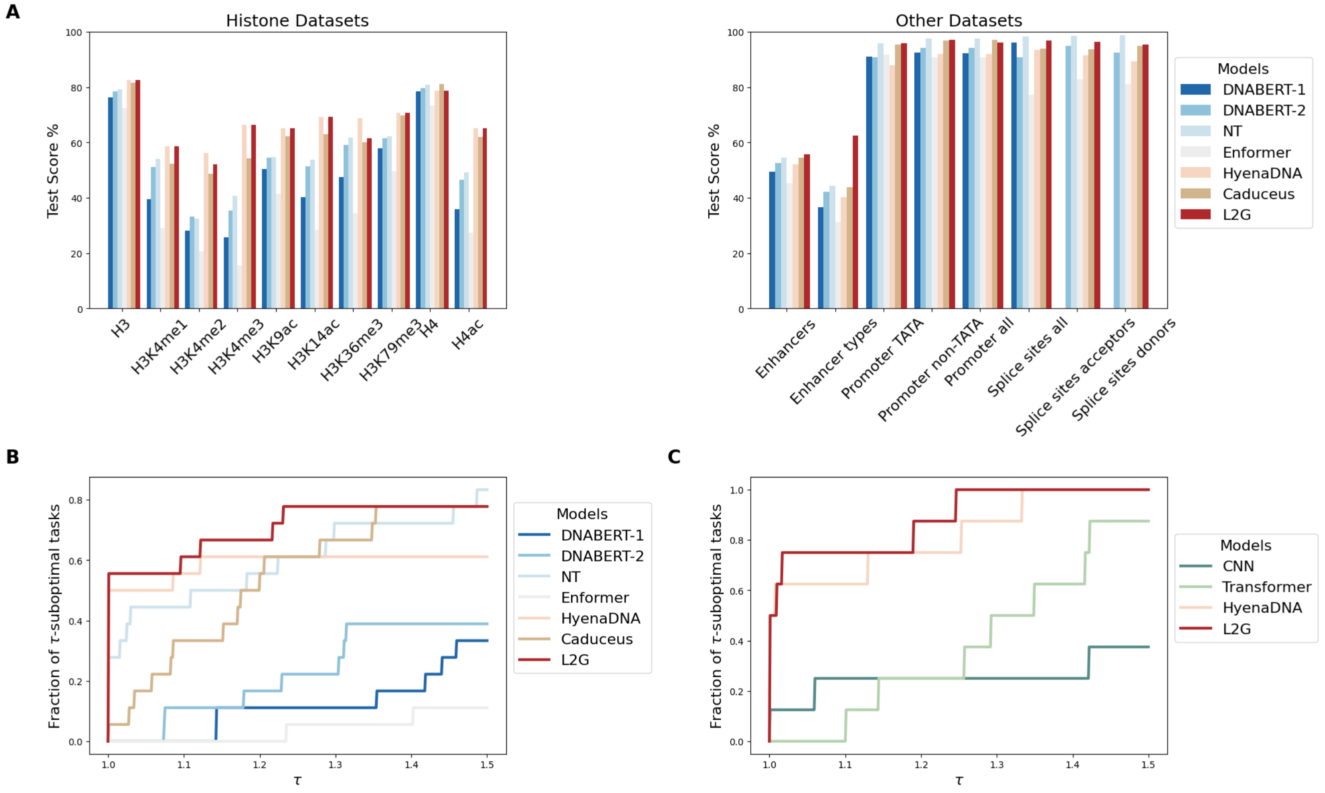


**Figure S1:** L2G matches or outperforms recent DNA foundation models on the Nucleotide Transformer (NT) Benchmarks and Genomic Benchmarks. **A**. Test scores on the Nucleotide Transformer Benchmarks for histone mark prediction tasks (left) and enhancer, promoter, and splice site prediction tasks (right). The bar for DNA-BERT-1 is missing because it could not predict splice sites. **B**. Aggregating results on the Nucleotide Transformer Benchmarks (**Table** S4) using performance profiles [10]. Larger values (fractions of tasks on which a method is within a $\tau$-factor of the best) indicate better performance. L2G's curve in the top-left corner demonstrates it is often the best or second best method. **C**. Aggregating results on the Genomic Benchmarks, which include supervised CNN and transformer baselines.

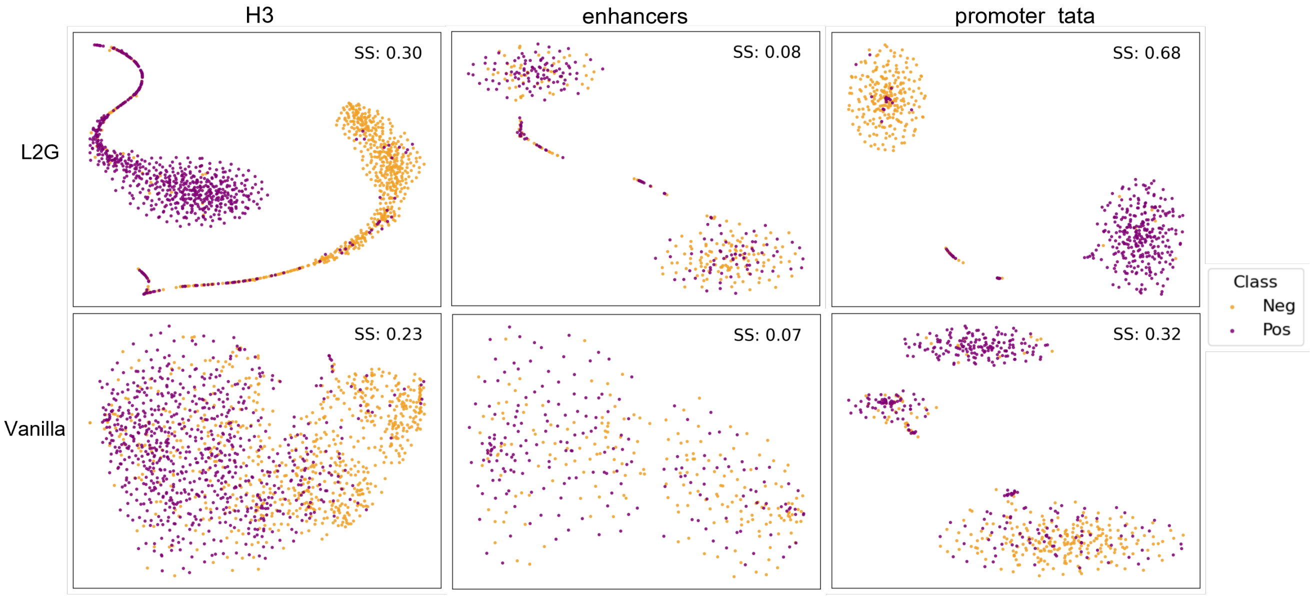


**Figure S2:** Visualization of the learned embedding of target modality data for models trained with L2G (top), and vanilla fine-tuning (bottom). The target modality data is from three representative downstream tasks from the NT Benchmark: histone modification (H3, left), enhancer regions (enhancers, middle), and promoter regions (promoter_tata, right). Pink dots represent positive class, while yellow dots indicate negative class. Each plot includes the Silhouette Score, a widely used metric for evaluating cluster separation. The score ranges from -1 to +1, where: +1 indicates well-separated clusters, 0 suggests indifferent clusters, and -1 indicates potential misclassification of points between clusters [11].

| Forward | Reverse | # Seqlets | Motif | Q value | Match |
|---|---|---|---|---|---|
| 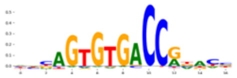 | 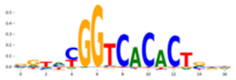 | 1059 | AP-1 | 0.006236 | 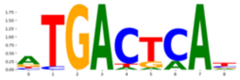 |
| 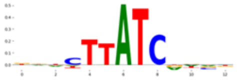 | 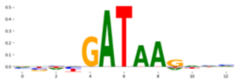 | 616 | GATA | 0.028951 | 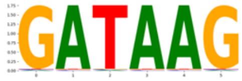 |
| 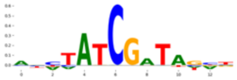 | 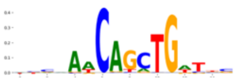 | 325 | da | 0.005764 | 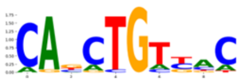 |
| 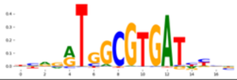 | 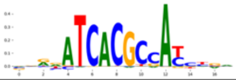 | 187 | SREBP | 0.001854 | 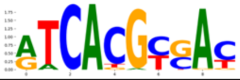 |

**Figure S3:** Motifs discovered for developmental enhancers by L2G. Details include forward and reverse sequences, seqlet count, motif name, Q-Value, and closest database match. The Q-Value is statistical measure that represents the false discovery rate (FDR) for the motif. Lower Q-values indicate more significant results.

| Forward | Reverse | # Seqlets | Motif | Q value | Match |
|---|---|---|---|---|---|
| 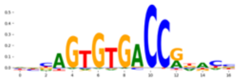 | 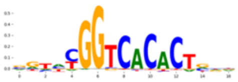 | 848 | Ohler1 | 0.000003 | 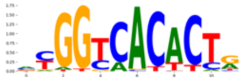 |
| 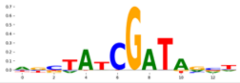 | 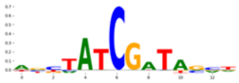 | 704 | DRE (Dref) | 0.043078 | 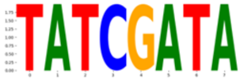 |
| 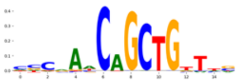 | 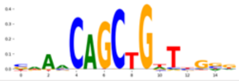 | 263 | crp | 0.013335 | 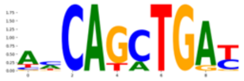 |
| 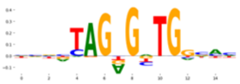 | 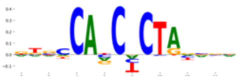 | 247 | Ohler7 | 0.051141 | 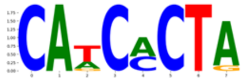 |
| 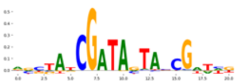 | 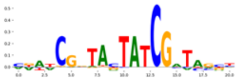 | 63 | BEAF-32 | 0.021631 | 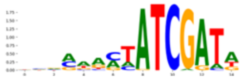 |
| 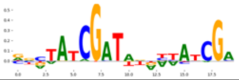 | 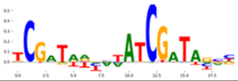 | 30 | DRE+DRE | 0.019005 | 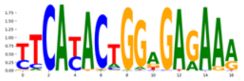 |

**Figure S4:** Motifs discovered for housekeeping enhancers by L2G. Details include forward and reverse sequences, seqlet count, motif name, Q-Value, and closest database match.
